# Transcriptomic profiling of a novel gastric implantation model identifies mechanisms and pathways that drive implantation into explanted human peritoneum

**DOI:** 10.1101/2025.04.14.648628

**Authors:** Deanna Ng, Karineh Kazazian, Kiera Lee, Yi Qing Lu, Aiman Ali, Karina Pacholczyk, Savtaj Brar, James Conner, Igor Jurisca, Christopher Allan McCulloch, Dae-Kyum Kim, Carol Jane Swallow, Marco Magalhaes

**Affiliations:** Department of Surgery, University of Toronto; Lunenfeld-Tanenbaum Research Institute, Sinai Health System; Department of Oral and Maxillofacial Sciences, Sunnybrook Health Sciences Centre; Faculty of Dentistry, University of Toronto; Pathology and Laboratory Medicine, Sinai Health System; Osteoarthritis Research Program at Schoeder Arthritis Institute and Data Science Discovery Centre for Chronic Diseases at Krembil Research Institute, UHN; and Departments of Medical Biophysics and Computer Science, University of Toronto; Division of Thoracic and Upper Gastrointestinal Surgery, Department of Surgery, Faculty of Medicine and Health Sciences, McGill University; Cancer Research Program, Research Institute of McGill University Health Centre and Montreal General Hospital Foundation, McGill University Health Centre

**Author notes:** Joint last author (Carol Jane Swallow, Marco Magalhaes).

## Abstract

Epithelial cancers such as stomach and ovary cancer tend to metastasize to the peritoneum, often leading to intractable disease and poor survival. Currently, the mechanisms that enable gastric cancer cells to penetrate the mesothelium, and to implant, invade, and survive in the peritoneal niche are poorly understood. To investigate these mechanisms, we developed a novel human peritoneal explant model. Briefly, fresh peritoneal tissue samples from abdominal surgery patients were cultured on top of a layer of GFP-labeled human gastric adenocarcinoma cells (AGS); 2% of these cells implanted into the peritoneum. The transcriptomic profile of the implanted AGS cells was compared to the profile of AGS cells that failed to implant using RNA sequencing. Differentially expressed genes were enriched significantly (fold change>2) for genes that enable cell adhesion, motility, and membrane depolarization. We compared this list of genes with a previously identified peritoneal metastasis whole exome sequencing dataset (SRP043661). Upon further analysis based on subcellular localization, cell adhesion and cytoskeletal organization, we found nine core “peritoneal implantation” genes. We functionally validated these genes with CRISPR knockout and assessed peritoneal implantation and invasion using the human peritoneal explant model described above. From these data we identified ADAM12 as a key player of peritoneal metastasis. Knock out of *ADAM12* significantly impaired peritoneal metastasis *in vivo and ex vivo*. Exploration of three publicly available independent datasets indicated that *ADAM12* is indeed clinically relevant in peritoneal metastasis. To explore the role of ITGAβ1 in mediating cell-matrix interactions in the presence of ADAM12, we performed ITGAβ1 pull-down assays followed by mass spectrometry analysis in ADAM12 WT cells. ADAM12 KO cells show a marked disruption of the ITGAβ1 interactome in GCa cells. Key cytoskeletal proteins such as MYH14, MYH10, MYH9, ACTA2, SPTN1 and TPM1–TPM4 were found to be interactors with ITGAβ1 in ADAM12 WT cells but not ADAM12 KO cells. Our approach and the new data identify a distinct peritoneal metastasis gene set that facilitates implantation and invasion of gastric cancer cells within the peritoneum. Disruption of these pathways with peritoneal-directed therapies has the potential to improve survival in patients with high-risk primary gastric cancer.

## Introduction

Gastric cancer has a predilection for metastatic spread to the peritoneum, a phenomenon that is associated with intractable disease and poor survival^1–3^. The mechanisms that allow gastric cancer (GCa) cells to penetrate the mesothelium, implant, and survive in the peritoneal niche are poorly understood. Therefore, uncovering the mechanism underlying the interaction of GCa cells with the peritoneum is of critical importance. But current efforts to characterize this mechanism are limited by an absence of tools that enable study of real-time *in vivo* cancer cell invasion, and a lack of high-fidelity models of the human peritoneum. Existing models of peritoneal metastasis largely include (i) *in vitro* models, (ii) *in vivo* animal models, and (iii) artificial human peritoneal tissue, including biomimetic three-dimensional cell cultures and organ-on-a-chip devices ^4,5^.

Despite their utility, traditional cell culture methods and mouse models often fail to capture the intricate characteristics of tumors and the dynamic microenvironment observed in human gastric cancer^6^. These models lack essential features such as native stromal architecture, immune components, and mesothelial surfaces, limiting their translational relevance. While ex vivo models preserve the native tumor microenvironment^7,8^ and enable real-time observation of tumor dynamics^8^, many existing studies are limited by their narrow focus on late-stage disease or isolated gene pathways, providing only a limited view of the metastatic process. Key interactions between tumor cells and their microenvironment, including signaling pathways involving stromal cells, immune cells, and extracellular matrix components, remain underexplored^9,10^.

RNA sequencing offers a powerful, unbiased approach to interrogate the molecular mechanisms driving peritoneal dissemination. Unlike targeted methods, RNA-seq enables comprehensive transcriptomic profiling, capturing global gene expression changes associated with tumor progression, invasion, and microenvironmental interactions. Prior studies have identified expression signatures linked to peritoneal relapse and dissemination, yet most rely on samples from primary tumors or fully established metastases, missing the critical early events that initiate implantation^11–13^.

To address this gap, we developed a human peritoneal explant model that preserves the native tissue architecture and allows for unbiased high-resolution analysis of early-stage tumor cell adhesion and invasion. When coupled with RNA-sequencing, this system enables discovery of the gene networks and signaling pathways activated during the initial steps of peritoneal metastasis, providing a platform for identifying novel biomarkers and therapeutic targets with greater clinical relevance.

## Results

### Implanted (IMP) and Non-IMP cells cluster distinctly and are involved in cell adhesion, ECM organization

A key question in peritoneal metastasis is understanding the molecular differences between gastric cancer cells that implant into peritoneal tissue versus those that do not implant. Accordingly, we sought to identify differentially expressed genes (DEGs) that drive peritoneal adhesion and implantation.

In this approach for selecting ECM adhesion and implantation, GFP-labelled AGS gastric cancer cells were seeded on a 10 cm plate and overlayed with fresh human peritoneum placed on top of the cells (i.e., mesothelial surface facing AGS cells). After 48h, most AGS cells (98%) remained in a monolayer attached to the plate, but 2% of AGS cells associated with the peritoneal tissue sample. The tissue sample was incubated with trypsin + collagenase for 30 min, washed and cells were FAC sorted to yield an Implanted (IMP) and Non-Implanted (Non-IMP) AGS cell sample for RNA sequencing (Fig. 1a).

**Figure 1.**
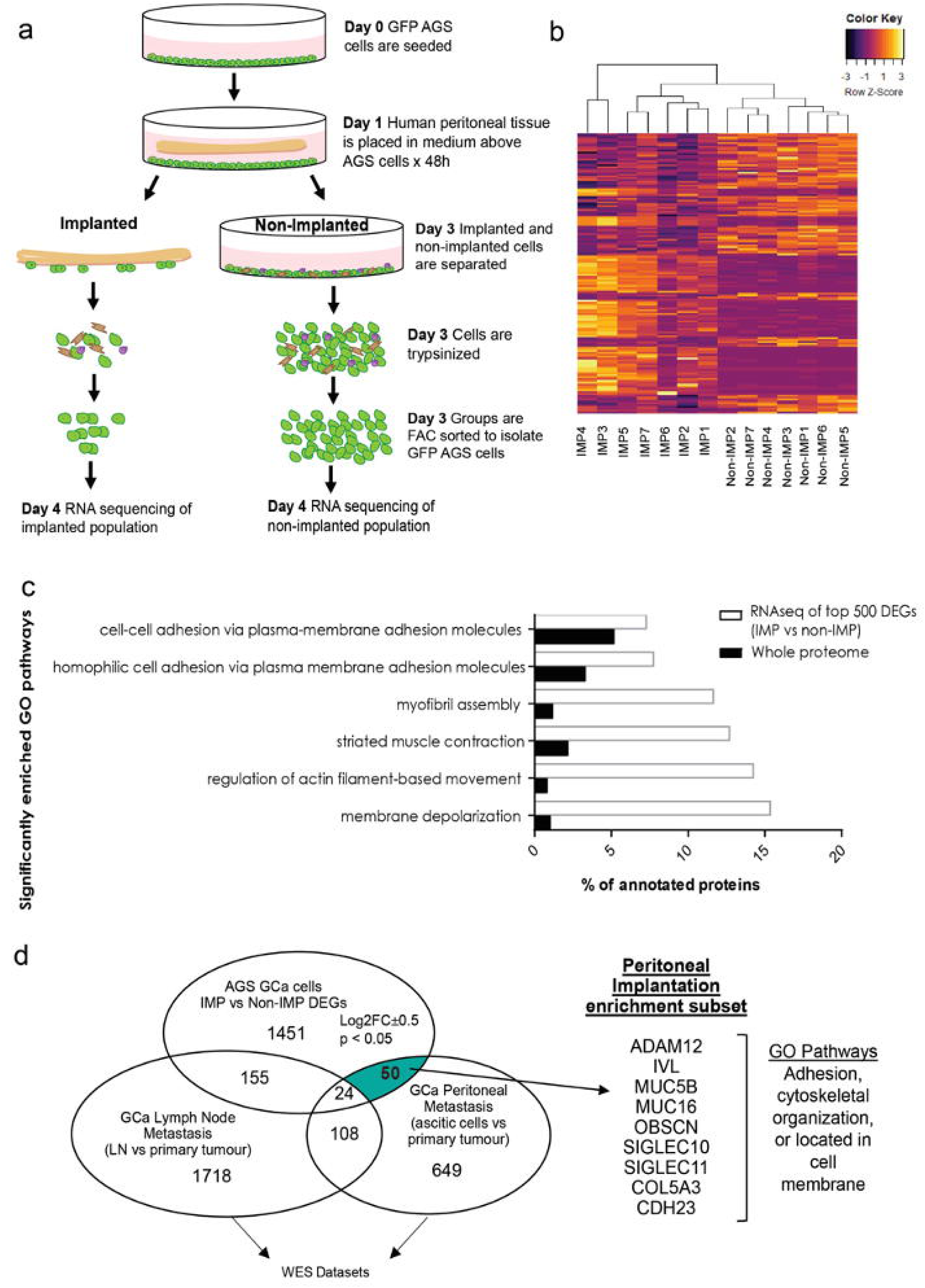
Investigation of genes driving implantation using Ex Vivo peritoneal model. a) Experimental setup. GFP labelled AGS gastric cancer cells are seeded on 10 cm plate. The next day, fresh human peritoneum from the operating room is placed in medium above the cells, mesothelial side down. After 48 h, the majority of AGS cells (98%) remain in a monolayer on the plate, but 2% of AGS cells are associated with the peritoneal tissue sample. This was incubated with trypsin + collagenase x30 min, then washed and cells FAC sorted to yield and implanted (IMP) AGS cell sample for RNA sequencing. The Non-Implanted (Non-IMP) cell sample was treated the same. b) Heat map showing unsupervised clustering of top 150 differentially expressed genes. c) Gene Ontology analysis of top 500 upregulated Differentially Expressed Genes (DEGs) in IMP vs Non-IMP demonstrates that pathways related to cell adhesion, cytoskeletal organization and membrane depolarization are significantly upregulated (p<0.05). d) Venn diagram showing overlap of genes in three datasets: 1. IMP vs Non-IMP dataset; 2. Peritoneal metastasis (PM) WES dataset SRP043661 (ascitic cells vs primary tumour); 3. Lymph node metastasis WES dataset SRP093339 (LN vs primary tumour). 50 genes (Peritoneal implantation subset) were included in both IMP dataset and the PM dataset, but excluded from the LN dataset (Table 1). Genes were triaged by subcellular localization (cell membrane) and GO function (adhesion, cytoskeletal organization) to derive a peritoneal implantation enrichment subset (9 genes). Validation by RT PCR on cell extracts yield a final candidate list of 5 genes (IVL, MUC16, MUC5B, ADAM12, OBSCN).

**Table 1.** Peritoneal implantation subset (50 genes)

|  |  |  |  |  |
| --- | --- | --- | --- | --- |
| IRF2BP2 | NTM | LRMP | PPFIA2 | EPB41L3 |
| DIAPH3 | IFI16 | FAM186A | ELL | NFAT5 |
| EGLN3 | MUC16 | FERMT2 | SIGLEC10 | KLK2 |
| GCDH | OBSCN | PIK3R5 | SPAG17 | SIGLEC11 |
| ZNF678 | MUC5B | IGFN1 | ARRDC4 | AIPL1 |
| SPATA6 | CDH23 | MYO3A | CLCA2 | FLT3 |
| KIAA0430 | PAPPA2 | SPTA1 | NR4A1 | IVL |
| CCDC62 | KNDC1 | SORCS3 | AKR1C2 | ADAM12 |
| BCO2 | KLF17 | SLC6A17 | IPO7 | PXDN |
| LINGO4 | COL5A3 | SYT5 | DAGLA | ANKRD45 |

While searching for differential expression, 257 genes were down-regulated (by < 0.5 fold change) and 1423 genes were up-regulated (by >2-fold change; all analyses were P < 0.05). Principal component analysis (PCA) showed distinct clustering of samples characterised by the implantation into peritoneal tissue (Sup Fig.1). The top 150 differentially expressed genes also formed distinct groups based on unsupervised clustering (Fig. 1b). To further investigate the function of DEGs in implantation to peritoneal tissue, upregulated DEGs in IMP vs Non-IMP were examined with Gene Ontology analyses using Geneontology.org. These analyses demonstrated that pathways related to cell adhesion, cytoskeletal organization and membrane depolarization were upregulated (Fig. 1C; P < 0.05).

A second key question was whether the genes differentially expressed in peritoneal implantation were uniquely involved in peritoneal metastasis rather than being associated with other metastatic pathways, such as lymph node metastasis. This distinction is crucial, as peritoneal metastases arise through distinct mechanisms compared to lymphatic spread. To address this distinction, we compared the overlap of genes in three datasets: 1. our IMP vs Non-IMP dataset; 2. Peritoneal metastasis (PM) WES dataset SRP043661 (ascitic cells vs primary tumour)^14^; 3. Lymph node (LN) metastasis WES dataset SRP093339 (LN vs primary tumour) ^15^. In the peritoneal metastasis dataset, whole-exome sequencing was performed, which compared primary tumors to malignant ascites samples from eight patients with peritoneal carcinomatosis^14^. In the lymph node metastasis dataset, whole-exome sequencing was performed, which compared primary tumours to metastatic lymph node samples from 5 patients^15^. We found that 50 genes (peritoneal implantation subset) were included in both IMP vs Non-IMP dataset and the PM WES dataset but were excluded from the LN WES dataset (Fig. 1d and Table 1).

To refine our understanding of the specific molecular drivers of peritoneal implantation, we further triaged genes based on subcellular localization and Gene Ontology (GO) function This is critical as peritoneal metastasis follows a unique dissemination pathway that requires tumor cells to detach, survive in ascitic fluid, and adhere to the mesothelial lining before invading underlying tissues. Accordingly, we sought to identify genes with direct functional relevance to these processes rather than those broadly involved in metastasis. Genes were triaged by subcellular localization (cell membrane) and GO function (adhesion, cytoskeletal organization) to derive a peritoneal implantation enrichment subset (9 genes): CDH23, MYO3A, SIGLEC10, SIGLEC1, MUC5B, MUC16, IVL, OBSCN, ADAM12. Validation by qRT-PCR of cell extracts from 3 additional independent peritoneal assays as described above yielded a final candidate list of 5 genes (IVL, MUC16, MUC5B, ADAM12, OBSCN) (Supp Fig. 2).

**Figure 2.**
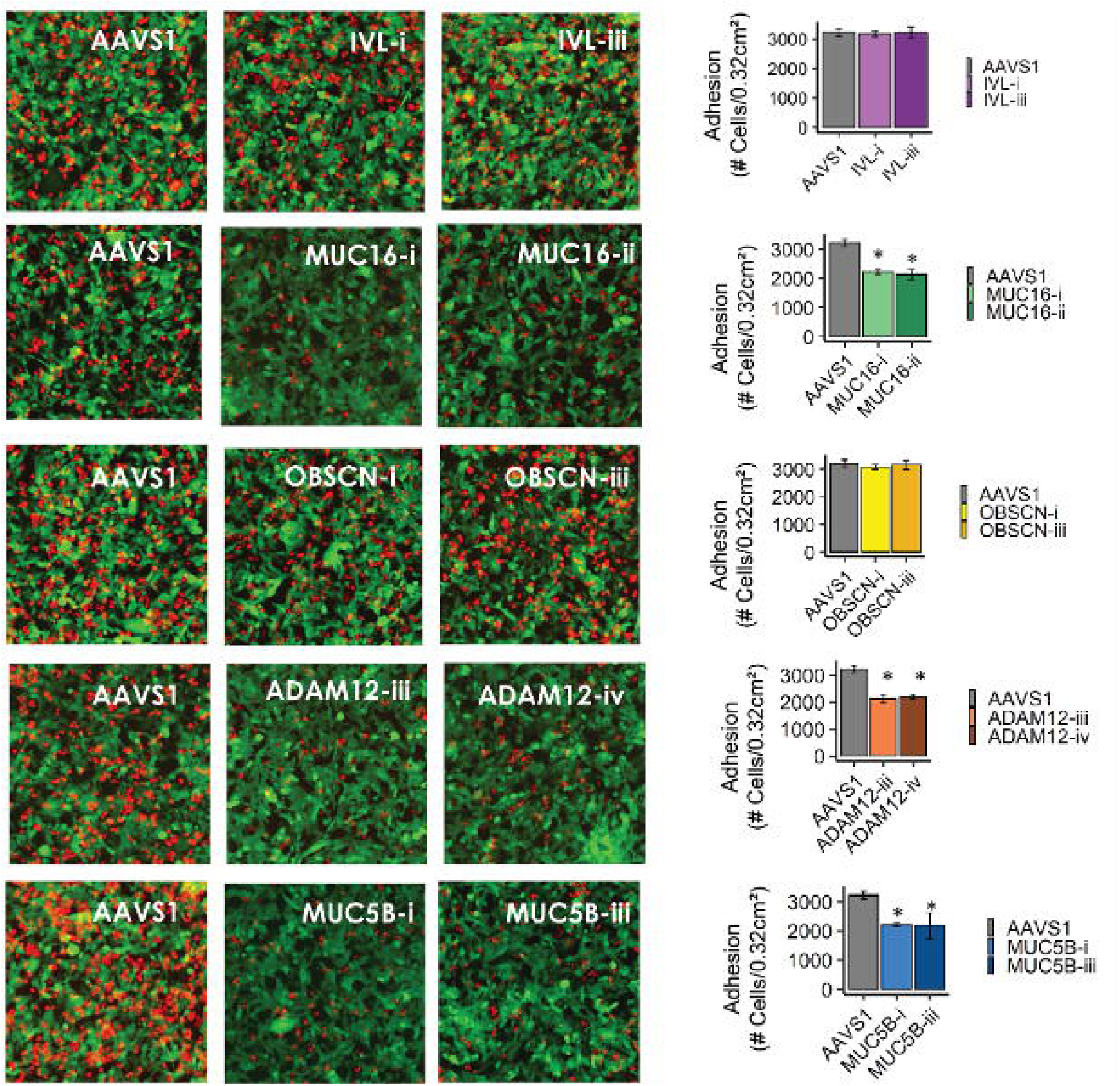
Mesothelial adhesion is decreased in AGS gastric cancer cells with ADAM12, MUC16 and MUC5B depletion but not in AGS gastric cancer cells with OBSCN and IVL depletion. In a Mesothelial Adhesion Assay, AGS gastric cancer cells express mCherry while the mesothelial cells express green fluorescent protein (GFP). mCherry-AGS gastric cancer cells attach to GFP expressing mesothelial monolayer after 30 minutes. En-face fluorescence time-lapse images of mesothelial adhesion assay showing presence of AGS cells adherent to the mesothelial layer. Quantification of cells over time shows reduced adhesion of cells with ADAM12, MUC16 and MUC5B depletion. Data are presented as mean ± SEM of 3 independent experiments, *=p<0.001 vs AAVS1 control.

### ADAM12 facilitates mesothelial adhesion in vitro

A critical initial step in peritoneal metastasis is the adhesion of cancer cells to the mesothelial lining of the peritoneum, a process that facilitates subsequent invasion and tumor progression. To investigate the role of ADAM12 and other candidate genes in this process, we performed an in vitro mesothelial adhesion assay using GFP-expressing mesothelial cells and red-stained AGS gastric cancer cells. This assay allowed for direct visualization and quantification of cancer cell attachment to the mesothelial monolayer. Compared to AAVS1 control cells, depletion of MUC16, ADAM12, and MUC5B significantly reduced the number of cancer cells adhering to the mesothelial layer, suggesting their role in promoting peritoneal implantation (Fig. 2). In contrast, depletion of OBSCN and IVL did not alter adhesion, indicating that these genes may not be directly involved in the early adhesive interactions required for peritoneal dissemination.

### Depletion of ADAM12 suppresses implantation and invasion

To determine whether ADAM12 and MUC16 contribute to gastric cancer implantation and invasion, we first assessed their role in an ex vivo peritoneal implantation assay. Using a previously established model^8^, we observed that AAVS1 control cells and MUC5B-depleted AGS cells implanted within the peritoneum were proliferating by day 3 while implantation in ADAM12 and MUC16-depleted AGS cells was markedly suppressed (Fig. 3a). Further, depletion of ADAM12 markedly suppressed the invasion of AGS cells over the course of 3 days (Fig. 3b).

**Figure 3.**
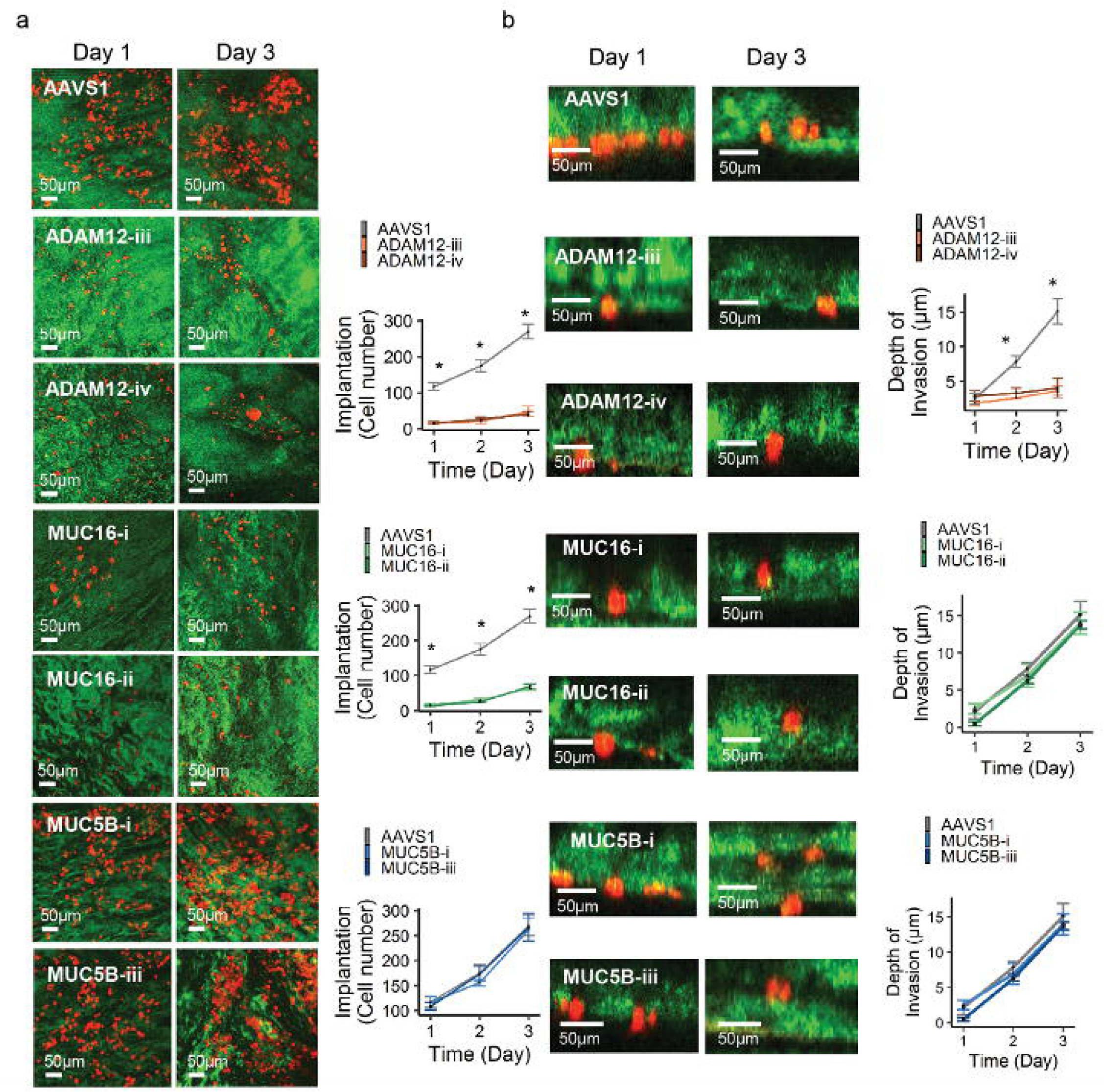
Implantation and invasion into ex vivo peritoneal tissue is decreased in AGS gastric cancer cells with ADAM12, MUC16 but not MUC5B depletion. a) En-face fluorescence images of ex vivo peritoneum shows that implantation of AGS gastric cancer cells is decreased with depletion of ADAM12 or MUC16, but not MUC5B. Quantification of cells over time shows reduced implantation of cells with ADAM12, MUC16 but not MUC5B depletion. Data are presented as mean ± SEM of 3 independent experiments, *=p<0.001 vs AAVS1 control. b) Fluorescence images of ex vivo peritoneal assay showing invasion of AGS gastric cancer cells is decreased with depletion of ADAM12 but not MUC5B or MUC16. Quantification of depth of invasion over time shows reduced invasion of cells with ADAM12 but not MUC5B or MUC16 depletion. Data are presented as mean ± SEM of 3 independent experiments, *=p<0.001 vs AAVS1 control.

To further investigate the potential role of ADAM12 and MUC16 as drivers of aggressive behavior in gastric cancer cells, we conducted functional studies to evaluate the impact of gene depletion on key metastatic traits such as motility and invasion. Following gene depletion, cell motility was assessed using wound-healing (scratch) assays in which the rate of gap closure provided a measure of the cells’ migratory capacity. Directional migration was reduced in AGS cells with depletion of ADAM12 or MUC16 (Fig 4a). To further evaluate invasive potential, Matrigel-coated Transwell invasion assays were performed which quantified the ability of the cells to degrade extracellular matrix components and traverse the membrane. ADAM12 and MUC16 depletion markedly suppressed the migration of AGS cells over 24 hours while only ADAM12-KO suppressed invasion over 36 hours (Fig. 4b).

**Figure 4.**
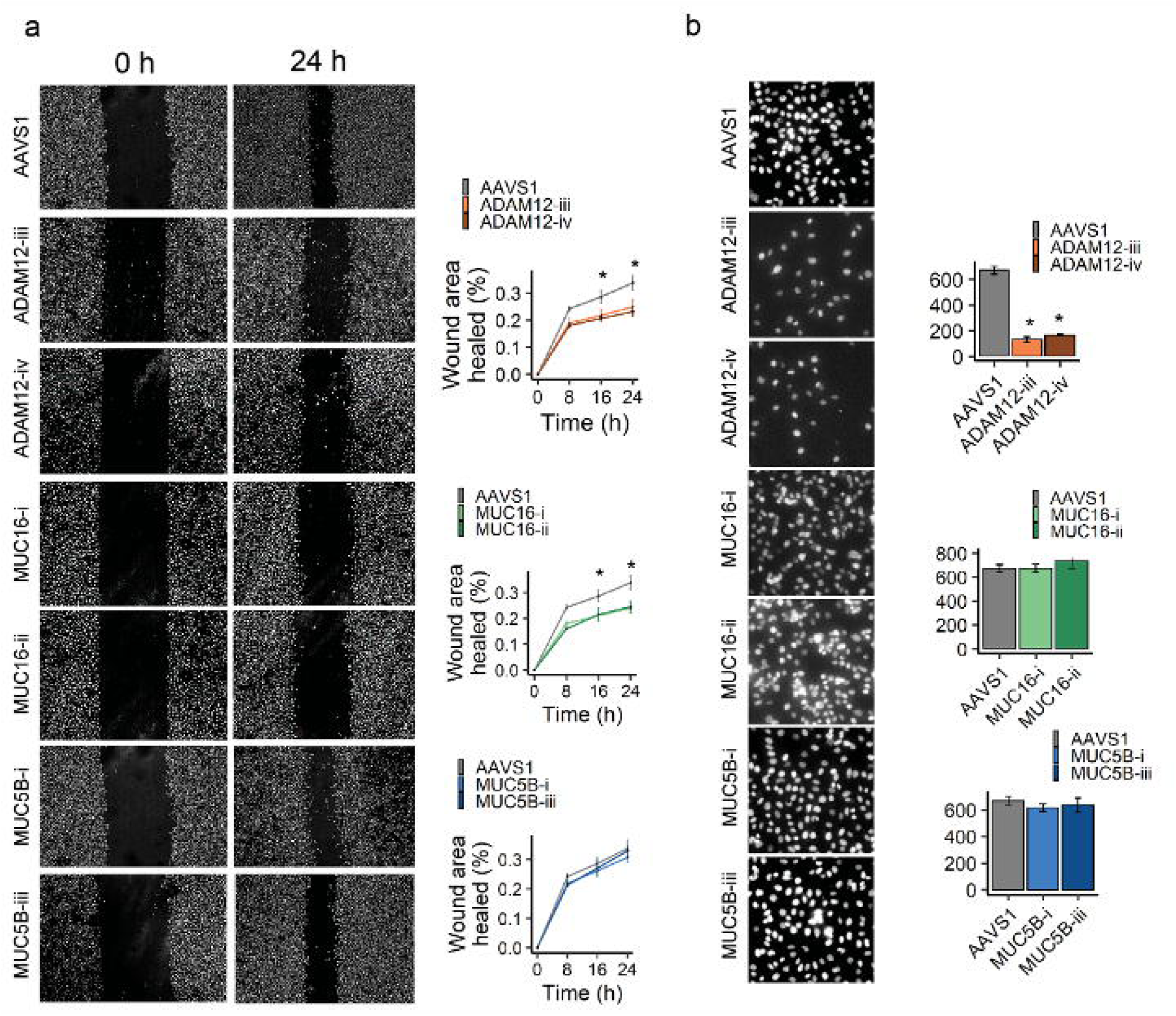
ADAM12 promotes directional migration and Transwell invasion by AGS gastric cancer cells. a) Directional migration, assessed in a scratch wound healing assay, is reduced in AGS cells with depletion of ADAM12 or MUC16 (Left). Summary of 3 independent experiments (right). Data are presented as mean ± SEM of 3 independent experiments, *=p< 0.001 vs AAVS1 control. b) Invasion, assessed in a Transwell invasion assay, is reduced in AGS cells depleted of ADAM12 but not MUC16 (Left). Summary of 3 independent experiments (right). Data are presented as mean ± SEM of 3 independent experiments, *=p< 0.001 vs AAVS1 control.

### ADAM12 facilitates peritoneal metastasis in vivo

To test the hypothesis that ADAM12 activity promotes GCa peritoneal metastasis, we employed a xenograft model in NOD SCID Gamma mice. MKN-45 cells were injected intraperitoneally into the right lower quadrant of NOD/SCID/Gamma mice and followed over 3 weeks. Three weeks following *i. p.* injection of MKN45 cells, the extent of peritoneal metastasis was described using the mouse Peritoneal Carcinomatosis Index (PCI) adapted from Biastiaenen 2020 et al., calculated by assessing 7 different regions of the peritoneal cavity.

We observed evidence of the tumor invading out of the peritoneum in control GFP-shRNA tumors but not in ADAM12-shRNA tumors (Fig. 5a). GFP shRNA tumors were larger and fixed to the underlying organs such as the diaphragm (Fig. 5b) and the small bowel mesentery (Fig. 5c) while ADAM12 shRNA tumours were more mobile and grossly less attached. H&E staining of the tumour nodules at the diaphragmatic peritoneal surface showed the presence of cancer invading the diaphragmatic muscle in GFP control but not ADAM12 shRNA tumours (Fig. 5b). H&E also showed invasion into mesentery in GFP-shRNA tumors, but not ADAM12-shRNA tumours (Fig. 5c). We quantified the total tumour weight of all tumour nodules harvested in the intra-abdominal cavity and found that tumour weight was decreased with ADAM12 depletion (Fig. 5d). ADAM12 depletion was also associated with a decreased mouse Peritoneal Carcinomatosis Index score (Fig. 5e). In all MKN45 xenografts, knockdown of ADAM12 in vivo to 25-45% of GFP control was confirmed (Fig. 5f).

**Figure 5.**
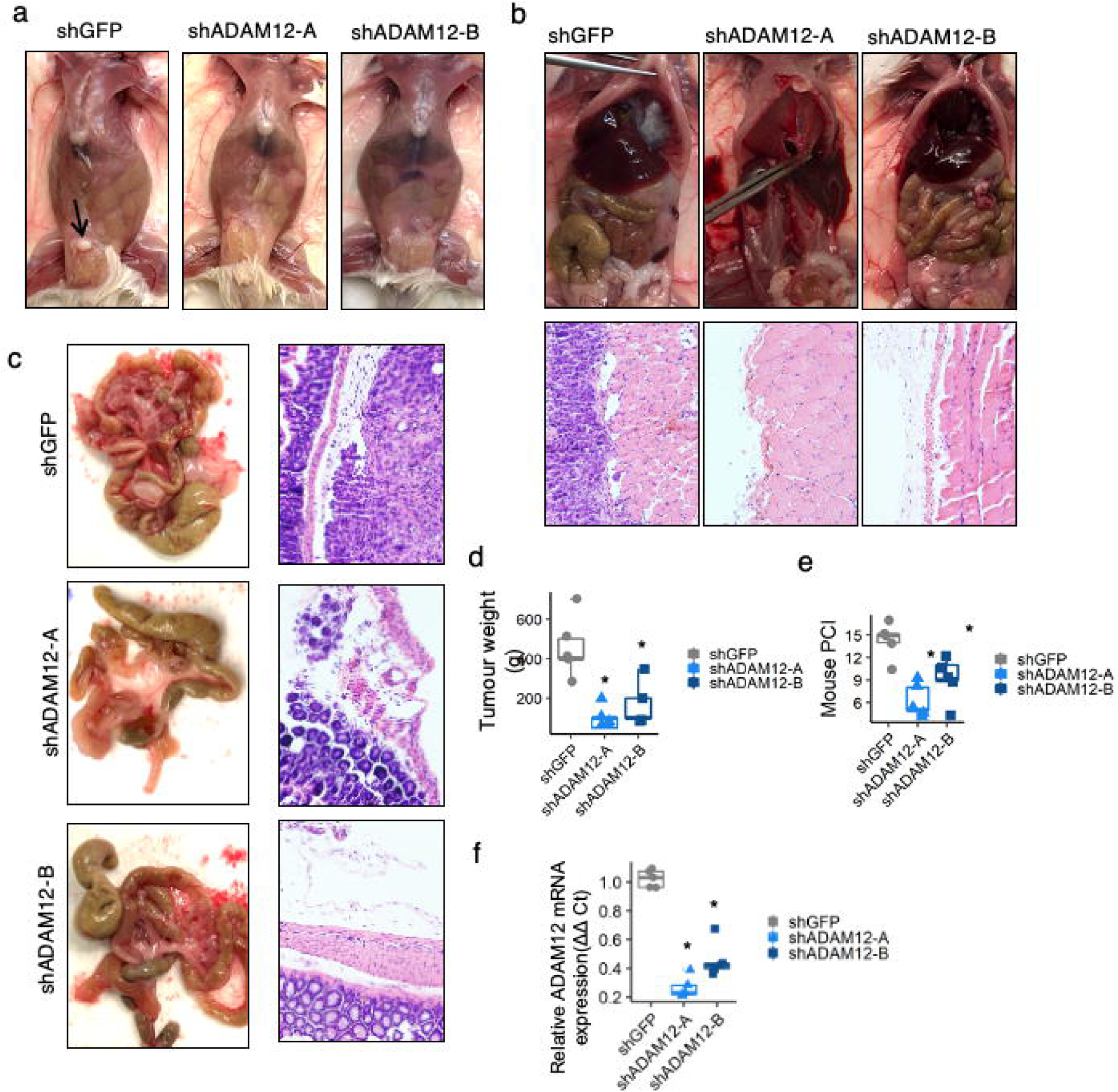
Requirement of ADAM12 for MKN-45 xenograft intraperitoneal metastasis. a) 1×10^6^ MKN-45 cells were injected intraperitoneally into the right lower quadrant of NOD/SCID/Gamma mice and followed over 3 weeks. a) Tumour invaded out of the peritoneum in GFP-shRNA tumours (arrow) but not in ADAM12-shRNA tumours. b) Intraperitoneal diaphragmatic tumour of xenografts showing modest suppression in ADAM12-vs. GFP-shRNA after 3 weeks. Representative images of formalin-fixed paraffin-embedded sections from diaphragm sections stained using H&E, showing invasion into underlying muscle in GFP-shRNA tumors, but not ADAM12-shRNA tumours. c) Tumour burden on small bowel showing modest suppression in ADAM12-vs. GFP-shRNA after 3 weeks. Representative images of formalin-fixed paraffin-embedded sections from small bowel sections stained using H&E, showing invasion into mesentery in GFP-shRNA tumors, but not ADAM12-shRNA tumours. d, e) Intraperitoneal tumour burden was assessed via mouse Peritoneal Carcinomatosis Index (PCI), adapted from Biastiaenen 2020 et al., with a maximum score of 21. Intraperitoneal tumour burden of xenografts showing modest suppression in ADAM12-vs. GFP-shRNA after 3 weeks, quantified by total tumour weight (d) and mouse PCI score (3); *, P < 0.01 versus ADAM12 shRNA. f) ADAM12 knockdowns in tumours persisted over 3 weeks.

### ADAM12 is dysregulated in advanced gastric cancer

A key question we sought to address was whether ADAM12 expression is clinically relevant in gastric cancer, particularly in advanced disease and peritoneal metastasis. To investigate this, we analyzed publicly available transcriptomic and genomic datasets to assess ADAM12 expression patterns, mutational status, and association with clinical outcomes in gastric cancer.

Using publicly available datasets, we found that ADAM12 is implicated clinically as it is dysregulated in advanced gastric cancer. In the TCGA dataset of 315 patients, ADAM12 expression is increased in tumor vs. normal mucosa (Fig. 6a). Meanwhile, ADAM12 is mutated or amplified in nearly 10% of esophagogastric carcinomas and cancers (Fig. 6b), along with a wide variety of cancer types, including lung, skin, and colorectal cancers (Fig. 6c). In the GSE5459 dataset, genome-wide mRNA expression profiles of 200 primary gastric tumors from the Singapore patient cohort was obtained. Here, we found that overall survival was reduced in gastric cancer patients with increased ADAM12 expression (divided by median expression) (Fig. 6d). In addition, increased ADAM12 expression was associated with a higher stage, where patients with Stage 3 and 4 disease had a higher expression of ADAM12 compared to patients with Stage 1 and 2 disease (Fig. 6e). In the ACRG Gastric cohort (GSE62254, n=305), microarray profiles from 300 gastric tumors from gastric cancer patients was provided. In this cohort, in patients with Stage 2 or 3 diseases, the development of Peritoneal Metastasis was associated with a higher level of ADAM12 expression in primary tumors compared to patients with no peritoneal metastasis (Fig. 6f). Finally, Tanaka et al. (GSE162214) comprehensively analyzed cancer cells purified from malignant ascitic fluid samples and their corresponding cell lines from 98 patients; increased ADAM12 expression was seen in ascites compared to cell lines derived from primary tumor (Fig. 6g).

**Figure 6.**
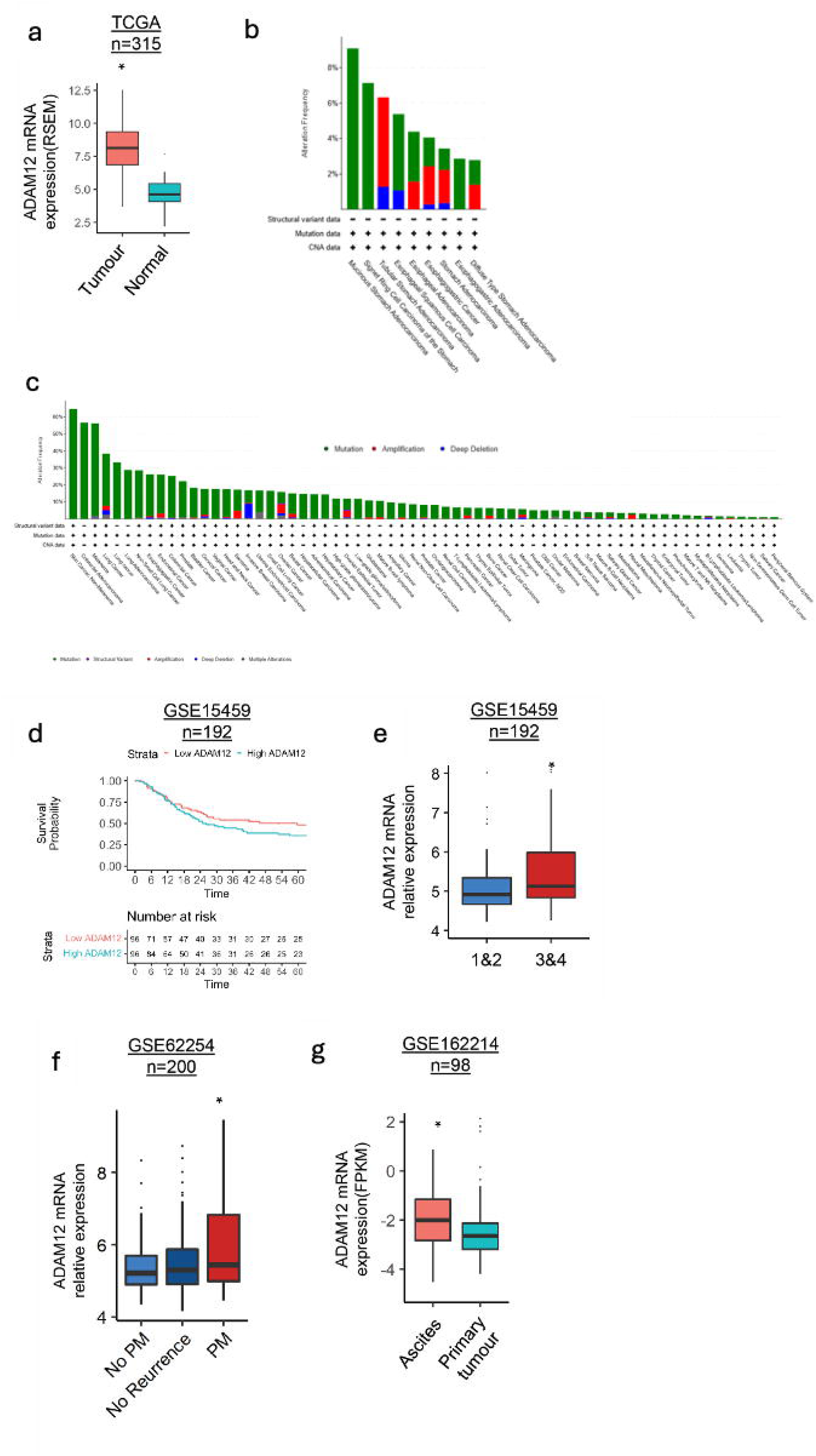
ADAM12 is dysregulated in advanced gastric cancer. a) ADAM 12 expression is increased in tumour vs. normal mucosa in 315 patients who underwent resection of primary gastric cancer. Data are derived from the TCGA dataset. bc) cBioPortal OncoPrint displaying the genetic alteration of ADAM12 in various b) esophagogastric and c) other cancer types. d) Overall survival is reduced in gastric cancer patients with increased ADAM 12 expression in 192 patients who underwent resection of primary gastric cancer, GSE15459. e) Increased ADAM12 expression is associated with higher stage (* vs Stage 1/2, p<0.05), GSE15459 (n=192). f) Development of Peritoneal Metastasis (PM) is associated with a higher level of ADAM12 expression than No PM in patients with Stage 2 or 3 disease (*p<0.05) GSE62254 (n=305). g) Increased ADAM12 expression is seen in ascites compared to cell lines derived from primary tumour, (*p<0.05), GSE166214 (n=98).

### ADAM12 interacts with *ITGA****β****1*to facilitate peritoneal metastasis

Given that ADAM12 is a known interactor of ITGAβ1, and that integrins orchestrate dynamic signaling during cell migration and tissue invasion, we hypothesized that ADAM12 may modulate or stabilize the β1 integrin interactome to facilitate metastatic progression. To explore the role of ITGAβ1 in mediating cell-matrix interactions in the presence of ADAM12, we performed ITGAβ1 pull-down assays followed by mass spectrometry analysis of ADAM12 WT cells. ADAM12 KO cells show a marked disruption of the ITGAβ1 interactome in GCa cells. Key cytoskeletal proteins such as MYH14, MYH10, MYH9, ACTA2, SPTN1 and TPM1–TPM4 were identified as interactors with ITGAβ1 in ADAM12 WT cells but not in ADAM12 KO cells (Fig. 7a). The high proportion of these interacting proteins involved in the pathways seen in only ADAM12 WT cells but not in ADAM 12 KO cells reinforces the hypothesis that ITGAβ1 (a high abundance receptor for collagen) is a key mediator of cytoskeletal dynamics and metastatic behavior, particularly in ADAM12-mediated mechanisms. This interactome analysis provides insight into the molecular players involved in ITGAβ1-mediated signaling and highlights their functional relevance in cell migration, invasion, and metastasis.

**Figure 7.**
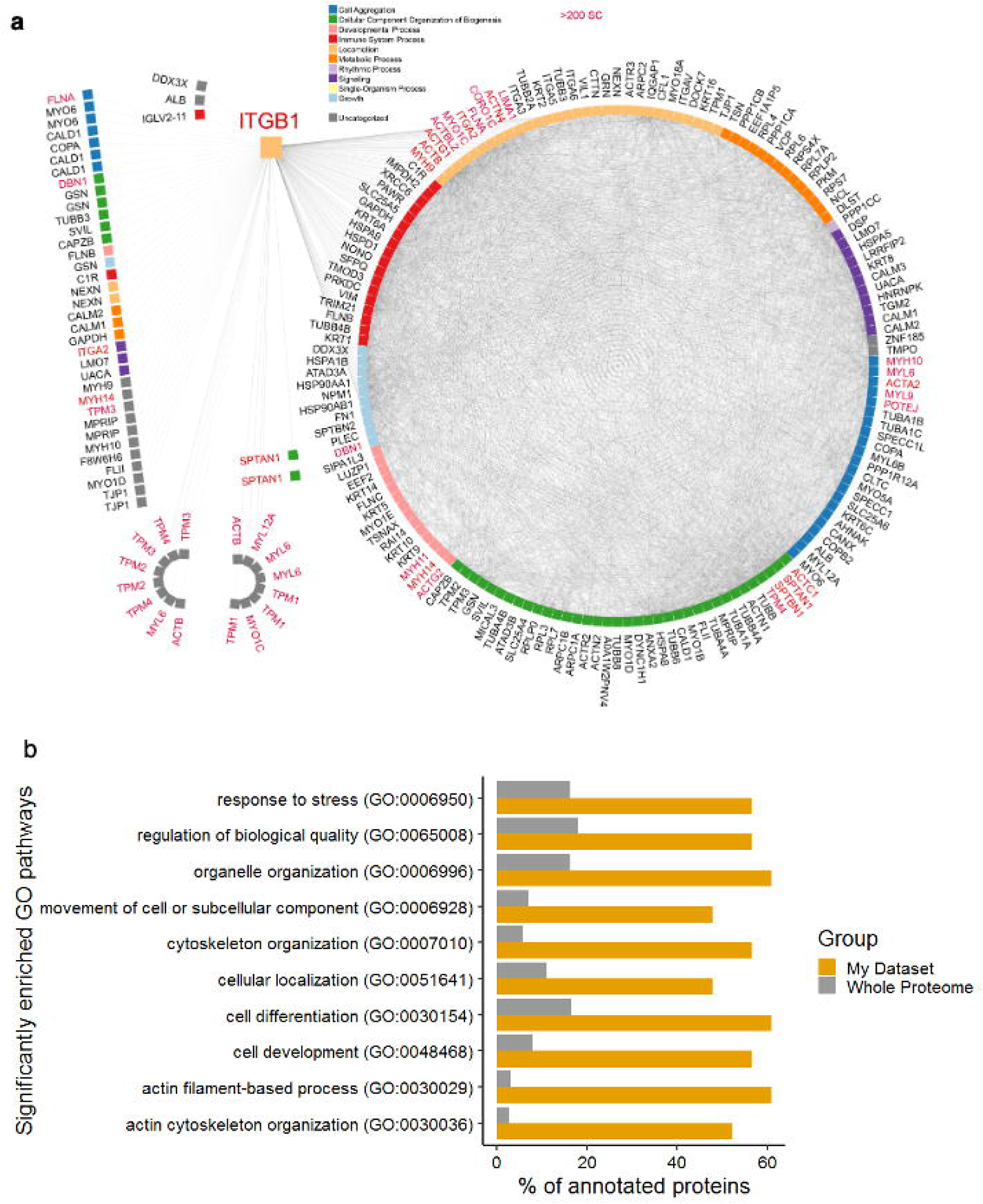
**ITGA**β**1 Interactome in ADAM12 WT Cells.** a) ITGAβ1 pull-down assays followed by mass spectrometry analysis in ADAM12 WT vs KO cells identified a network of over 200 proteins in ADAM12 WT cells but not ADAM12 KO cells interacting with ITGAβ1, many of which are involved in processes critical for cytoskeletal organization, cell adhesion, and motility. Proteins identified were categorized based on their associated Gene Ontology (GO) biological processes, represented by color coding in the figure. Prominent biological processes include cytoskeletal organization (cyan), actin filament dynamics (red), cellular localization (purple), and locomotion (blue), reflecting the central role of ITGAβ1 in regulating cellular movement and structural integrity. Proteins highlighted in red represent those with spectral counts (SC) > 200, indicating a strong interaction with ITGAβ1. b) Bar chart displaying the percentage of proteins annotated for specific GO biological processes in the dataset (orange) versus the whole proteome (gray). The GO terms are ranked based on their relevance to the dataset, focusing on processes such as actin cytoskeleton organization (GO:0030036), actin filament-based processes (GO:0030029), cytoskeleton organization (GO:0007010), and movement of cell or subcellular component (GO:0006928). Pathways with significantly higher enrichment in the dataset compared to the whole proteome indicate a strong association of the identified proteins with cellular motility, cytoskeletal organization, and stress response.

The central role of these proteins was also seen in Gene Ontology (GO) enrichment analysis in which the proteins that interacted with ADAM12 (in ADAM 12 WT cells) and ITGAβ1 were overrepresented (adjusted Pval<0.05) in pathways related to cytoskeletal organization and cellular motility (Fig. 7b). The most enriched GO term was actin cytoskeleton organization: nearly 60% of proteins in the dataset associated with actin. Similarly, actin filament-based processes and cytoskeleton organization were highly enriched, emphasizing the dataset’s focus on structural components of the cell involved in migration and adhesion. Collectively, these findings support the hypothesis that proteins identified in the dataset are deeply involved in pathways central to cytoskeletal remodeling, cell migration, and stress response, which are critical for the progression and metastasis of cancer. This aligns with the role of key proteins such as ITGAβ1 and ADAM12 in mediating these processes.

## Discussion

In this study, using a novel *ex vivo* model of human peritoneal tissue, we systematically identified genes involved in the initial attachment process of peritoneal metastasis. Fresh human peritoneal tissue samples were obtained and suspended, mesothelial layer down, above a monolayer of GFP-labeled AGS human gastric cancer that had been seeded. Through FACS and RNA sequencing, we compared the genomic profile of AGS cells that successfully implanted into the peritoneum compared to those that failed to implant. The results of this model were validated with *ex vivo*, *in vitro,* and *in vivo* experiments.

Through differential gene expression analysis, the *ex vivo model* implicated *MUC5B*, *MUC16,* and *ADAM12* in peritoneal metastasis. None of these proteins were previously known to be involved in this process. However, only ADAM12 was implicated in implantation and invasion *ex vivo*. Notably, although the depletion of *MUC16* and *ADAM12* decreased implantation, they did not eliminate peritoneal metastasis completely. The primacy of ADAM12 in our analyses of peritoneal metastasis is consistent with its role in the proliferation, invasion, and metastasis of a variety of cancers in the peritoneal cavity as well as small cell lung cancer and non-melanoma skin cancers. Further, these data support the hypothesis that peritoneal implantation and invasion involves a complex interplay of additional genes, outside of matrix metalloproteinases and mucins^16,17^.

While depletion of MUC16 impaired implantation, invasion was unaffected. This finding can be attributed to the properties of *MUC16* as an adhesion molecule that binds to mesothelin on the mesothelial cells. Hence, depletion of MUC16 impairs the ability of gastric cancer cells to implant onto the peritoneum. However, in the very small number of cells that overcame impaired adhesion, invasion into the peritoneum was unaltered. This was reaffirmed in our in vitro experiments in which MUC16 did not affect Transwell invasion. Interestingly, MUC5B depletion did not affect GCa implantation, migration, and invasion *ex vivo* and *in vitro.* This finding contradicts a study by Lahdaoui et al. (give reference here) in which MUC5B was found to be involved in GCa migration and invasion *in vitro*. A potential explanation for this discrepancy is that their study used KATOIII cells that strongly express MUC5B compared with AGS cells that exhibit low expression of MUC5B expression^18^.

Several potential explanations may account for how *ADAM12* is involved in implantation. First, ADAM12-L can bind to ITGA-B1 on the mesothelial cells. Next, ADAM12-S can interact with syndecan and ITGA-B1, promoting the formation of adhesion complexes ^19^. Hence, depletion of ADAM12 results in decreased mesothelial adhesion, which leads to decreased implantation. Further, the steps of invasion are altered due to the proteolytic activity of ADAM12 which mediates cleavage of Insulin-like growth factor binding proteins or through its role in ectodomain shedding, activating membrane-anchored proteins, such as sonic hedgehog or EGFR. This notion is consistent with the published literature in which expression of *ADAM12* (but not *MUC16)* was found to be increased in patients with peritoneal metastasis, compared to other forms of metastasis. *MUC16* is typically found in ovarian cancer peritoneal metastasis; however, this was not found to be significant in gastric cancer, possibly due to the low expression of *MUC16* in gastric cancer. ADAM12 involvement *in vivo* was further confirmed where knockdown of *ADAM12* resulted in decreased peritoneal implantation. Our study is the first to demonstrate that *ADAM12* is implicated in peritoneal metastasis.

The strengths of our study include the novel use of an *ex vivo* model, which enabled the identification of genes involved in the initial steps of peritoneal metastasis. Indeed, in our model, the cancer cells are only in contact with the peritoneal tissue for 48 hours. This approach confers an advantage as other forms of differential expression analysis performed between matched primary tumor and peritoneal metastasis often use peritoneal metastasis samples that have been established for months. As a result, they do not capture the initial steps of adhesion and invasion. In addition, commonly utilized peritoneal metastasis samples often become fibrotic. Hence, bulk sequencing may not inaccurately reflect the actual mechanisms occurring.

The limitations of our study are first, the tissue was left suspended over the cancer cells for 2 days. As gastric cancer tumor adhesion is estimated to occur over a span of 30 minutes, we were not able to restrict the study to either the process of adhesion or invasion. This limitation created challenges in delineating the specific genes involved in the peritoneal metastatic cascade. However, this instrument bias is unlikely to have affected our main conclusions as the steps of mesothelial adhesion, migration, and invasion were characterized in *in vitro* experiments. Further, the data from these experiments were confirmed in the *ex vivo model*. In addition, we are aware that we did not distinguish whether ADAM12-L or ADAM12-S was the key driver involved in peritoneal metastasis. This uncertainty is unlikely to have clinical consequences in the context of inhibition of general ADAM12 activity.

Collectively, these experiments have revealed a distinct peritoneal metastasis gene set that facilitates implantation and invasion of gastric cancer cells within the peritoneum. More importantly, we identified and functionally validated *ADAM12* as a potential gene of interest that regulates peritoneal metastasis by gastric cancer cells. Notably, there are currently available ADAM12 inhibitors which have been used *in vivo* that result in decreased ADAM12 proteolytic activity. By interrupting these pathways, peritoneal-directed therapies have the potential to improve the quality and length of life in patients with high-risk primary gastric cancer.

## Methods

### Cell culture

Cells were grown at 37°C in RPMI1640 (AGS, MKN-45) or DMEM (HEK293) and supplemented with 10% FBS (Wisent). Cell lines were obtained from ATCC in 2018, and were maintained between passages 4 and 15. HEK293 lines were a kind gift from the Daniel Schramek laboratory (obtained in 2020 and maintained between passages 8 and 28; Lunenfeld Tanenbaum Research Institute, Sinai Health System, Toronto, Canada). Cell lines were not further authenticated in our laboratory.

### Generation of stable cell lines (knockdown and overexpression)

MKN45 cell lines with stable knockdown of ADAM12 or control (GFP) were generated through lentiviral infection, using 2 individual short hairpin RNA (shRNA) constructs (Sigma). Lentiviruses were produced as described [74], and were used to infect the cells for 24h, followed by puromycin-(3μg/ml for MKN45 cells) mediated drug selection for 7 days. Confirmation of knockdown of ADAM12 was performed by RT-PCR with Taqman ADAM12 primers.

### Generation of stable cell lines (CRISPR KO)

Plasmid vectors: The lentiCRISPRv2 vectors were also provided as a gift from Jason Moffat, University of Toronto, Toronto, Canada. The pMD2.G (AddGene #12259), pCMV-VSV-G (AddGene #8454), and psPAX2 (AddGene #12260) vectors were obtained from AddGene, Watertown, MA, USA.

Ligation of vectors with gRNAs and Transformation: gRNAs were selected for each of the genes chosen for validation (selected from the TKOv3 pooled gRNA library used for the genome-wide screens). For each gene chosen for validation, 4 guides were chosen based on the highest guide score for that gene. Adeno-Associated Virus Integration Site 1 (AAVS1) was chosen as a positive control gRNA as it targets a noncoding region of the human genome and therefore should have no effect on the cell, Forward and reverse oligos for each chosen gRNA were mixed with T4 PNK and T4 ligation buffer (New England Biolabs, Ipswich Massachusetts, USA) and annealed in a thermocycler at 37°C for 30 minutes, then 95°C for 5 minutes, and then ramped down to 25°C at 5°C per minute. The pLCKO vector was digested with BsmBI first, then CIP (New England Biolabs, Ipswich Massachusetts, USA) was added and further incubated. The reaction was run on a 1% gel and the fragment was excised from the gel, following the kit manual for the QIAquick Gel Extraction Kit for each gel slice. Gels were made using UltraPure Agarose (Invitrogen, Thermo Fisher Scientific, Carlsbad, California, USA) and SYBR Safe DNA Gel Stain (Invitrogen, Thermo Fisher Scientific, Carlsbad, California, USA). The digested vector was combined with the annealed oligos and T4 ligase. The ligated vector was transformed into Stbl3 competent cells (Thermo Fisher Scientific), SOC-treated for 30 minutes, and plated on LB plates (Multicell, Wisent Inc., St-Bruno, QC) with 100 µg/ml of Ampicillin. After 24 hours, individual colonies were picked and grown in 5 ml of LB broth with 100-125 µg/ml of Ampicillin for 14-16 hours. The bacterial growth was pelleted via centrifugation and underwent a miniprep following the kit manual of the PureLink Quick Plasmid Miniprep Kit (Invitrogen, Thermo Fisher Scientific, Carlsbad, California, USA).

Generation of stable cell lines: AGS cell lines with stable knockout or control (AAVS1) were generated through lentiviral infection. Lentiviruses were produced as described [74], and were used to infect the cells for 24h, followed by puromycin-(2μg/ml for AGS) mediated drug selection for 7 days.

Confirmation of knock out: Confirmation of knock out was performed using Western Blots, RT-PCR, or Synthego ICE (https://ice.synthego.com/<u>#/</u>) as described). CHOPCHOP (https://chopchop.cbu.uib.no/) was used to select primers used for Sanger sequencing of each gene validated. The primers used are shown in the table below:

### RNA extraction and Real-time PCR

RNA was extracted from cell pellets using the RNeasy Mini Kit (Qiagen). RNA was quantified using NanoDrop ND-100 spectrophotometer, and 800-1600ng of RNA from each sample was used for reverse transcription. Samples were treated with RNase-free DNase (Invitrogen), and reverse transcribed with SuperScript II Reverse Transcriptase (18064-014; Invitrogen) using Random Primers (48190-011; Invitrogen) according to manufacturer’s instructions. Real-time RT-PCR was performed using Taqman PCR Master Mix (ThermoFisher) or SYBR Green PCR Master Mix (ThermoFisher) on an ABI 7900HT apparatus. Quantifications were normalized to control endogenous GAPDH. Data generated by PCR software (SDS2.2.2; Applied Biosystems) were analyzed using the 2−ΔΔCt method (24).

### Wound healing migration

1×10^6^ cells were seeded into 6 well plates and serum-starved for 18h before retrieval by scratch. Scratches were made manually using a P10 pipette tip, and migration was assayed for 24h using the INCell6000 analyzer equipped with a motorized stage and a live cell apparatus (37°C heated and humidified chamber with 5% CO2) with a 6-well plate adaptor. Data acquisition was performed continuously over the indicated time courses. Images were collected with a 10x objective lens. All hardware and image capture conditions were made possible, and images were analyzed, using MATLAB 9.4 2018.

Image analysis was carried out by measuring the total wound area in three fields per condition, at t=0, 8, 16, and 24 h. The residual wound area was expressed as a percent of original wound, which was itself highly reproducible, subtracted from 100 to yield the %healed at each time point, and compared across groups by t-test. The Bonferroni correction sets the significance at P<0.05 (two tailed).

### Transwell invasion assay

Transwell assays were performed as described previously (Oser et al., 2010). 8.0-mm Matrigel-coated Transwell supports from Becton Dickson Canada (BD) were used. 50,000 cells were resuspended in 500 µl 0.5% FBS/RPMI and plated in the upper chamber. The bottom chamber was filled with 1 ml 10% FBS/RPMI. Cells were allowed to invade for 24 h followed by fixation in 4% PFA. Intact membranes were stained with Alexa Fluor-488 Phalloidin (Life technologies) for 30 minutes and visualized in a Nikon Eclipse TE300 epifluorescence microscope. The cells at the upper side of the membrane were removed using a Q-tip. Cell invasion was calculated as the average cell number on the underside of the membrane compared with the control. Results were based on analysis of 20 fields in three independent experiments.

### Real time RT-PCR of tumour tissue

Mouse tissue was placed in Ambion RNAlater-ICE solution and thawed overnight at –20°C, then disrupted and homogenized in RLT buffer (RNeasy mini kit Qiagen) supplemented with β-mercaptoethanol using a rotor-stator homogenizer. The lysate was centrifuged at 4°C for 10 minutes at 14,000 rpm and the supernatant was used for RNA purification using the RNeasy mini kit following manufacturer’s protocol.

### Histology, immunochemistry and microscopy

Tissue samples were fixed in 10% formalin for 24h at room temperature. For human peritoneal tissue processing, serial coronal sectioning was performed at three levels, 50 mm apart. Each tissue edge was stained red with Tissue-Marking dye to orientate the specimen for paraffin embedding (Supp. Fig. 1A). The strips of peritoneal tissue were paraffin-embedded by lining the three strips parallel to each other, with the mesothelial layer oriented to one side (Supp. Fig. 1B). For mouse tissue, the small bowel, diaphragm, liver and lungs were processed for paraffin embedding. The tissue was cut into 20mm sections and was processed for paraffin embedding according to standard protocols. Sections (5 μm) were mounted onto slides and were stained with hematoxylin and eosin (H&E) according to standard protocols using the Veristain Gemini Automated Slide Stainer (Thermo Scientific) at the University of Toronto Dentistry Pathology Core, and examined with a Leica DMR upright microscope. For IHC staining, slides were stained with CDX2 or D2-40 antibodies using the fully automated Dako Omnis platform (deparaffinization and retrieval built-in), using the Dako OMNIS detection kit (Ref: GV800). Slides were exposed to high pH for 15 minutes, antibody incubation for 10 minutes, polymer detection for 20 minutes, substrate chromogen for 5 minutes, and finally Dako hematoxylin for 3 minutes. This was performed at the Department of Pathology at the Hospital for Sick Children, Toronto.

### Fluorescence-activated Cell Sorting

Cells were seeded at medium confluence in MatTek dishes and 10cm corning dishes and harvested the next day by trypsinization for flow cytometry-based live-death assay. Harvested cells were resuspended in flow cytometry sorting buffer (Hanks Balanced Salt Solution, 25 mM HEPES pH 7.0, 2 mM EDTA, 1% Fetal Bovine Serum) with 4′,6-diamidino-2-phenylindole (DAPI, 0.2 uM) at a concentration of 5 M cells per mL and were filtered by 40 μm nylon mesh to eliminate large aggregates. The stained samples were immediately analyzed on a Sony MA900 or Fortessa 20 flow cytometer. The data were subsequently processed with FlowJo software.

### RNA sequencing-Library prep and sequencing, RNA seq data analysis

RNA extractions from cell pellets obtained after FACS sorting were performed on the same day using RNeasy Mini columns (Qiagen). The sample RNA quality was accessed by Agilent Fragment Analyzer using High Sensitivity RNA Analysis Kit. A full-length cDNA library was first generated from high-quality total RNA using TaKaRa SMART-Seq v4 Ultra Low Input RNA Kit for Sequencing (Cat# 634890) according to manufacturer protocol. The quality and quantity of the purified full-length cDNA were measured by Agilent Fragment Analyzer and Qubit 2.0 (Thermo Fisher Scientific), respectively. The final sequencing libraries were prepared from full-length cDNA using Illumina Nextera XT Library Prep Kit according to manufacturer protocol. The cDNA libraries were checked with Agilent Fragment Analyzer for fragment size and quantified with qPCR using Collibri Library Quantification Kit (ThermoFisher) on a BioRad CFX96 Touch Real-Time PCR Detection System. Quality-checked libraries were then loaded onto an Illumina NextSeq 500 run with Illumina NextSeq 500/550 Hi Output Kit v2.5 (75 Cycles). Real-time base call (.bcl) files were converted to FASTQ files using Illumina bcl2fastq2 conversion software v2.17.The RNAseq reads were mapped to the human reference genome (Illumina iGenomes72 Homo sapiens UCSC hg38 build [88], annotations files from UCSC August 14-2015 release) using TopHat2 v2.1.1 [89].

Gene counts were obtained from featureCounts on the command line with options:-t exon-T 8 - s 2-g gene_id [90]. The table of raw counts was imported into R and converted to a DESeq2 object (DESeqDataSetFromMatrix using sample information) for processing, DESeq2 version 1.24.0 [91]. Genes with fewer than 10 counts across all samples were filtered out before differential expression analysis. Counts normalized by the DESeq2 rlog transformation were used for PCA and heatmaps of gene expression. Raw counts were used for differential expression analysis using the default parameters of the DESeq function. We called differential expression between Implanted cells (IMP) and Non-Implanted cells (Non-IMP) and applied a statistical cutoff (adjusted P < 0.1), to obtain refined sets of up-or down-regulated genes.

### TCGA and publicly available dataset analysis

TCGA data17 were obtained from the NIH National Cancer Institute GDC Data Portal Data release 8.0 (https://portal.gdc.cancer.gov/). OncoPrints of mutation frequencies were visualized using the cBioPortal platform as previously described (PMID: 38580884, 37668528, 22588877, 23550210).

Data was obtained from GEO GSE15459, GSE62254, and GSE15081. GSE15459 and GSE62254 were generated on the Affymetrix Human Genome U133 Plus 2.0 Array chip ^20,21^. Files were downloaded and preprocessed via RMA with the oligo package. The biomart package was used to annotate the files. The expression was then merged with the clinical data and differential analysis was performed with the DeSEQ2 package. GSE15081 was generated on a Hitachisoft AceGene Human Oligo Chip 30K 1 Chip Version. Raw data was downloaded and normalized for each channel, Cy3, and Cy5, independently ^12^. The Cy3 channel was loaded with control mRNA and Cy5 was loaded with experimental mRNA. The data was normalized by subtracting the mean of the channel and dividing by the standard deviation. The data was log-transformed and the change in expression of each probe in the experimental condition was calculated by subtracting Cy3 from Cy5.

### Integrin beta 1 immunoprecipitation

To prepare cell lysates for protein analysis, cells were lysed and lysate volumes were adjusted to ensure equal protein concentration across samples. 200µL of lysate was incubated with with 25µL of Dynabeads at 4°C to remove non-specifically binding proteins. ITGA-β1antibodies were crosslinked to beads by washing the beads with lysis buffer, adding 1µL of Rabbit ITGA-β1antibody or 5µL of non-specific IgG isotype to 100µL of lysis buffer, and incubating at room temperature for 40 minutes with agitation. Beads were washed 3 times with citrate phosphate buffer, twice with 0.2M triethanolamine solution, and incubated with 20mM DMP solution for 30 minutes at room temperature. Additional washes were performed with PBS/0.01% Tween 20. Beads were separated from the coating solution magnetically, washed with DPBS, resuspended in anrum-free medium, and stored at 4°C before use. Beads were added to AGS control and AGS ADAM12 KO cells and incubated for 3 h at 37°C. The cultures were gently washed with DPBS, lysed in CSKB buffer (containing 48 mM NaCl, 290 mM sucrose, 2.9 mM MgCl2, 9.65 mM PIPES, 1:50 diluted protease inhibitor cocktail, 1 mM PMSF, and 0.5% Triton X-100), and homogenized by vigorous vortexing. Beads were separated from the supernatant magnetically, washed 2X with 1 mL CSKB buffer, 3X with DPBS, and stored at −80°C before assaying with isobaric-label tandem mass tag mass spectrometry at the SPARC BioCentre (SickKids, Toronto).

### Statistical and computational biology analyses

Statistical analysis was done either by Student t-test with Bonferroni correction for continuous variables. Using the Kaplan–Meier method, OS and DSS curves were estimated, and then compared using the log-rank test. Analyses were performed using R version 3.5.1 software (Institute for Statistics and Mathematics of Wirtschaftsuniversität (WU), Wien, Austria: The R Project for Statistical Computing. https://www.rproject.org/). All p-values lower than 0.05 (two-sided) were considered statistically significant.

Physical protein-protein interactions for ITGA-β1 in ADAM12 WT cells were obtained from Integrated Interaction Database (IID; https://ophid.utoronto.ca/iid) ver.2021-05 ^22^. The network was visualized using NAViGaTOR v.3.0.19 ^23^. Exported SVG file was finalized with legend in Adobe Illustrator ver. 27.2.

## Acknowledgments

D.N. is funded by a Canadian Institutes of Health Research - Canada Graduate Scholarships Doctoral Award (CGS D). M.M. is funded by a Canadian Institutes of Health Research Project grant PJT 175128. The study is partially funded by the Mary Susan MacDonald Gastric Cancer fund to C.S.

