## Supplementary figures and images for "Transcriptomic profiling of a novel gastric implantation model identifies mechanisms and pathways that drive implantation into explanted human peritoneum"

### Supplemental 1

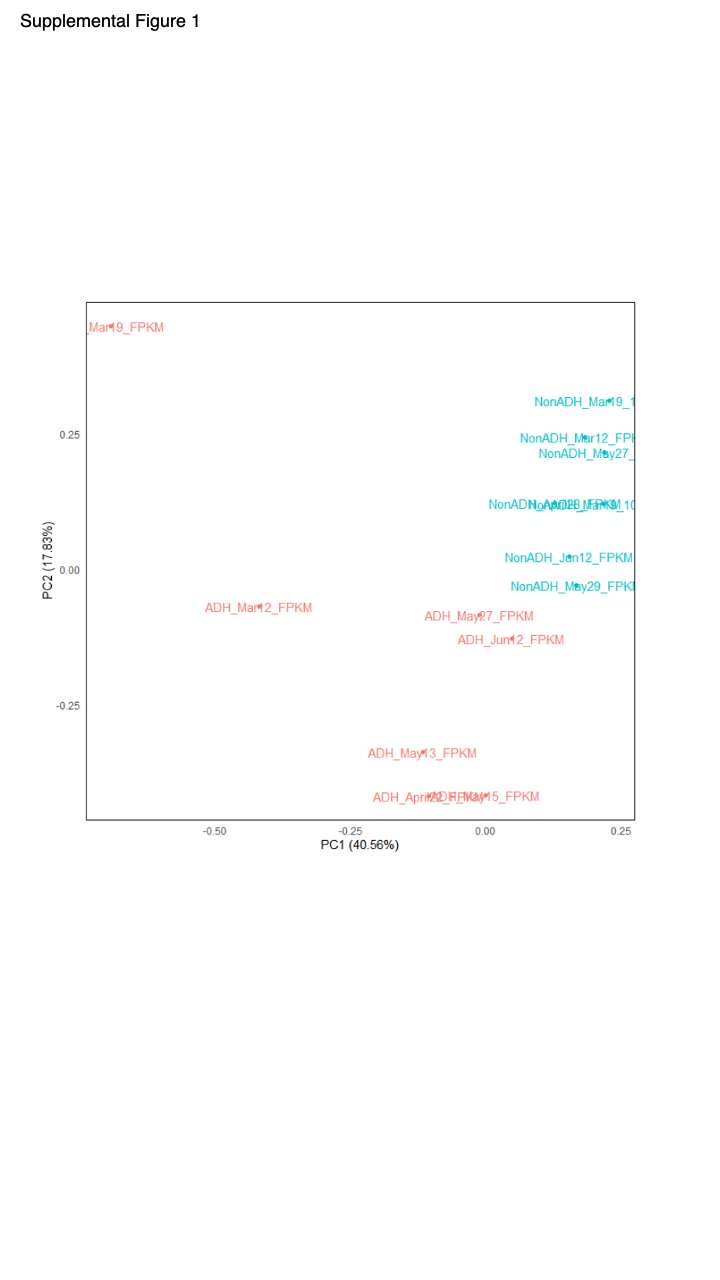

### Supplemental 2

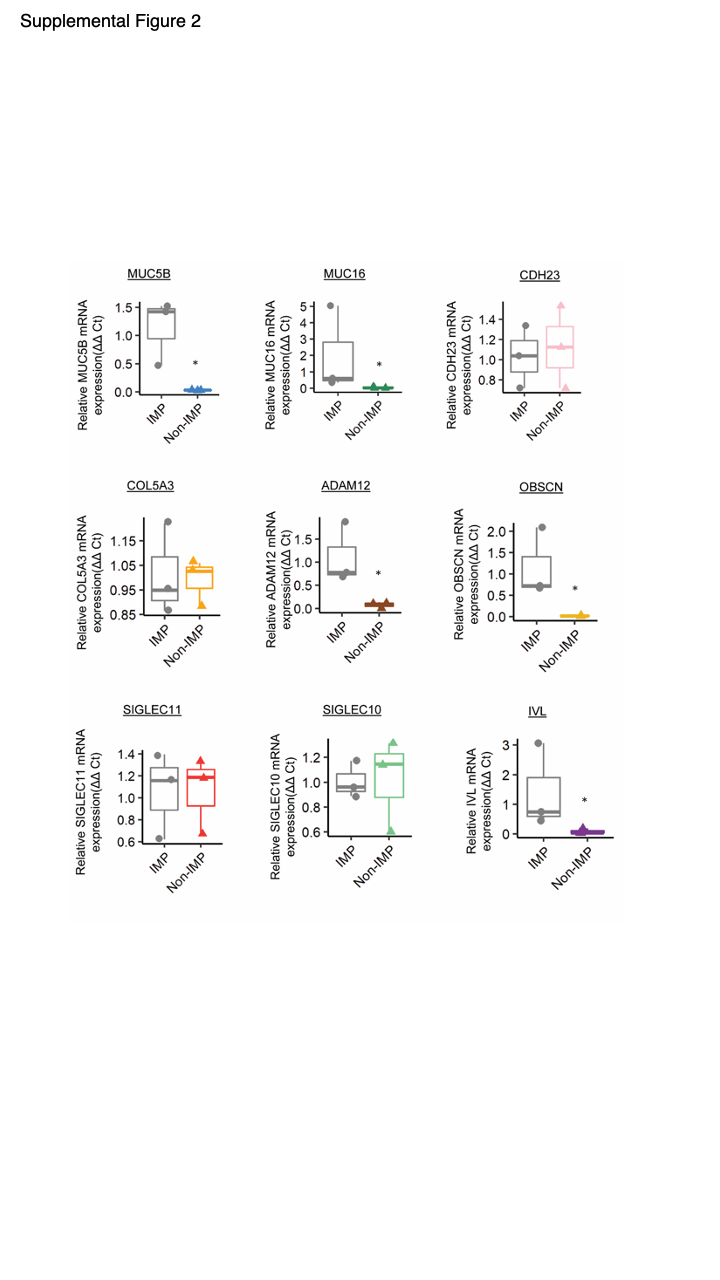
